# BioCPPNet: Automatic Bioacoustic Source Separation with Deep Neural Networks

**DOI:** 10.1101/2021.06.18.449016

**Authors:** Peter C Bermant

## Abstract

We introduce the Bioacoustic Cocktail Party Problem Network (BioCPPNet), a lightweight, modular, and robust UNet-based machine learning architecture optimized for bioacoustic source separation across diverse biological taxa. Employing learnable or handcrafted encoders, BioCPPNet operates directly on the raw acoustic mixture waveform containing overlapping vocalizations and separates the input waveform into estimates corresponding to the sources in the mixture. Predictions are compared to the reference ground truth waveforms by searching over the space of (output, target) source order permutations, and we train using an objective function motivated by perceptual audio quality. We apply BioCPPNet to several species with unique vocal behavior, including macaques, bottlenose dolphins, and Egyptian fruit bats, and we evaluate reconstruction quality of separated waveforms using the scale-invariant signal-to-distortion ratio (SI-SDR) and downstream identity classification accuracy. We consider mixtures with two or three concurrent conspecific vocalizers, and we examine separation performance in open and closed speaker scenarios. To our knowledge, this paper redefines the state-of-the-art in end-to-end single-channel bioacoustic source separation in a permutation-invariant regime across a heterogeneous set of non-human species. This study serves as a major step toward the deployment of bioacoustic source separation systems for processing substantial volumes of previously unusable data containing overlapping bioacoustic signals.

## Introduction

Bioacoustic source separation, informally referred to as the “cocktail party problem” (CPP), encompasses the problem of detecting, recognizing, and extracting meaningful information from conspecific signaling interactions in the presence of concurrent vocalizers in a noisy social environment^1^. Automatic source separation functions to isolate individual speaker vocalizations from a recording with overlapping signals. While human speech separation is an active and well-studied area of work involving the recent emergence of supervised deep neural networks (DNNs) for speaker separation^2–7^, the bioacoustic CPP remains comparatively understudied^8–11^. Occurrence of overlapping acoustic mixtures in recordings is especially common and problematic in the non-human bioacoustic domain. For instance, 58% of recordings of African elephant (*Loxodonta africana*) vocalizations contained at least two concurrent signalers^12^; 16% of sperm whale (*Physeter macrocephalus*) codas were overlapped by codas generated by another whale, and 22% were followed by another coda within 2s^13^, which is similar to how geladas (*Theropithecus gelada*) synchronize the onsets of their vocalizations^14^; in the NIPS4Bplus richly annotated birdsong dataset^15^, of the duration of recordings with vocalizations, nearly 20% contain simultaneously active classes. With no formal method for isolating individual-specific calls, biologists are often forced to omit data containing overlapping calls produced by simultaneously signaling vocalizers. In this paper, we offer a lightweight and modular neural network (NN) that acts on raw waveforms directly to reconstruct sources from recordings containing signal mixtures. Our machine learning (ML) model, the Bioacoustic Cocktail Party Problem Network (BioCPPNet), serves as an effective end-to-end source separation system to extract source estimations from composite mixtures, and we optimize the architecture specifically to accommodate bioacoustic vocal diversity across a range of taxonomic groups, even in limited data regimes.

Throughout the entire pipeline from data preprocessing to separator model evaluation, non-human bioacoustic source separation presents numerous challenges that complicate the problem relative to human speech separation. With non-human animals, technical and logistical difficulties associated with procuring bioacoustic data render it challenging to compile comparably-sized annotated datasets, and unique non-human vocal behavior demands differential treatment according to the species of interest. In the human speech domain, it is common practice to use large datasets of high-quality recordings downsampled to 8 kHz^2^ to minimize computational costs and to train models on segments several seconds in duration. However, this approach to data preprocessing is not always feasible for non-human animals. For instance, while the fundamental frequency (F0) of macaque vocalizations is on the order of 0.1-1 kHz^16^, which suggests that they could be treated similarly as human speech in terms of downsampling to reduce computational costs, the mean maximum F0 of bottlenose dolphin (*Tursiops truncatus*) signature whistles can range from 9.3-27.3 kHz^17^, which means that with sampling rates of 96 kHz, even the lowest order overtones approach or exceed the native Nyquist frequency. This is further exacerbated in animals such as bats, which can emit broadband calls with dominant frequencies ranging from 11-212 kHz^18^. Given the wide-ranging vocal behavior of non-human species, it is often infeasible to downsample recordings, resulting in long input sequences that pose a challenge to modern ML models. Though music source separation models such as Demucs^19^ have addressed higher sampling rates up to 44.1 kHz, the spectrotemporal parameters of non-human acoustic signals demand that we consider source separation with sampling rates up to 250kHz while still employing sufficiently long input sequences to reflect timescales related to the relevant biological behavior. The combination of relatively small datasets recorded with high sampling rates has significant implications for automatic bioacoustic source separation. In particular, time-domain separator NNs may struggle to capture long term temporal dependencies in the input time series, and recurrent-based approaches are often prohibitively expensive in terms of memory and computational costs. Further, heavyweight models such as those used to address human speech separation may overfit small datasets, impairing the models’ capacities to generalize. With BioCPPNet, we construct a lightweight model both to efficiently separate long mixture waveform sequences and to minimize the risk of overfitting small bioacoustic datasets.

In general, two families of algorithms have been implemented to solve the human speech CPP. These include frequency masking approaches^2,20^ that operate on handcrafted time-frequency representation (TFR) inputs generated by defining a mapping 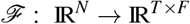 that projects the *N*-sample acoustic waveform to a spectral representation with *T* features of dimension *F*; and time-domain methods^3,5^ that operate on raw acoustic waveforms with learnable encoders. While existing research^21^ has compared various options for the TFR input, frequency masking-based methods for human speech separation commonly implement the mel-scaled short-time Fourier transform (STFT)-based spectrogram^22^ given its correspondence to human auditory processing and perception. In the bioacoustic regime, it remains unclear which TFRs may be optimal for ML applications. Though spectrograms or log-magnitude spectrograms are conventionally employed regardless of vocal characteristics, there exist numerous options for TFRs according to the vocal properties of the particular species of interest. For example, the Hilbert Huang transform (HHT) may provide advantages over Fourier-based analysis for transient signals such as those produced by sperm whales^23^; wavelet transforms have been used for automated birdsong detection^24^; though not extensively implemented in the bioacoustic literature, invariant scattering representations^25,26^ provide yet another option for representing animals sounds. Further, there also exist learnable spectral feature representations generated using convolutional neural networks (CNNs) acting on raw waveforms,^3,27,28^, but these fully learnable transforms remain unexplored in the bioacoustic domain. To address the fundamental bioacoustic representation problem, BioCPPNet provides a modular architecture to enable rapid experimentation using handcrafted or fully learnable encoders in combination with various fixed or learnable inverse transform decoders to recover the separated raw acoustic waveforms. In this study, we employ various STFT-based or learnable representations, but the architecture is compatible with other TFRs, which we defer to future studies.

Numerous additional factors further complicate the bioacoustic CPP. For instance, native environmental conditions often include acoustic interferences and energetic maskings that impair the ability of animals to recognize and discriminate signals. These interferences can consist of spectrotemporally overlapping calls produced by heterospecific signalers, other sources of biotic and abiotic noise, and anthropogenic sounds^1^. With these challenges, it remains difficult to construct sufficiently large and suitably annotated datasets to enable the training of supervised learning algorithms with access to ground truth signals. Given this limited and noisy data regime, we design BioCPPNet as a less complex model than many existing human speech separation networks^3–5^, and we integrate fixed or learnable noise reduction into the model architecture. Finally, compared to human speech separation applications, bioacoustic source separation models are more difficult to evaluate. While subjective assessment of the perceptual quality of human speech separation models can rely on panels of human listeners^3^, the human auditory system is often not well-suited for evaluating non-human vocalizations. This, in combination with the obscure interpretability of objective metrics such as the scale-invariant signal-to-distortion ratio (SI-SDR)^29^ in the context of different species with different vocal behaviors, implies that the evaluation of bioacoustic source separation models could benefit from other metrics. Qualitatively, we assess the separation performance of BioCPPNet using visual representations of the acoustic data in the form of spectrograms, and we quantitatively address the performance of BioCPPNet by considering downstream tasks. In particular, following the separation of the mixture into estimated source outputs, we feed the predictions into trained individual identity classifier models under the assumption that classification accuracy on downstream ML-based tasks serves as an objective proxy for the quality of reconstructed separated waveforms.

While the human speech separation problem is a competitive area of work, the bioacoustic CPP has received comparatively less attention, as current bioacoustic research often emphasizes other ML-based tasks such as automated detection and classification of bioacoustic sounds^30–32^. However, recent work has implemented both semi-classical and deep ML-based approaches to address bioacoustic source separation, employing time domain and TFR-based algorithms. Fast fixed-point independent component analysis (FastICA), principal component analysis (PCA), and non-negative matrix factorization (NMF) have been used to separate mixtures of overlapping frog sounds^9^. Mixed signals of overlapping dolphin signature whistles were separated with the joint approximate diagonalization of eigenmatrix (JADE) algorithm^8^. Time-frequency masking was used to address humpback whale (*Megaptera novaeangliae*) song separation^33^. More contemporary approaches have made use of DNNs, but they remain limited in their scope. For instance, bi-direction long short-term memory (BLSTM) networks were implemented to separate overlapping bat echolocation and communication calls^10^. In this application, the separation of composite mixtures containing signals of disparate classes or natures is more analogous to human music instrument separation into predefined classes^34^; it treats the separation problem as a multi-class regression problem and thus avoids the fundamental permutation problem^2^ of the CPP in which the source estimate channel order is arbitrary, potentially yielding conflicting gradients during training^35^. Other DNN-based approaches have adopted multi-step schemes to predict source number and to extract mask estimates using a segmentation-based approach to bat call separation optimized for fundamental frequency contours^11^; however, the omission of higher-order harmonics presents a dilemma since in biological systems, the inharmonic mistuning of spectral components may serve as an important acoustic cue for overcoming the CPP^36^; further, this bounding box-based approach performs best on signals containing low-to-moderate spectrotemporal overlap, which means they may be limited in their generalizability or practical implementation.

BioCPPNet represents a complete state-of-the-art permutation-invariant bioacoustic source separation pipeline across multiple species characterized by differential vocal behavior. The model is specifically designed to address key challenges of bioacoustic data, and we apply it to a diverse set of non-human species, including rhesus macaques (*Macaca mulatta*), bottlenose dolphins, and Egyptian fruit bats (*Rousettus aegyptiacus*). The multidisciplinary approach requires that we integrate a range of fields of study, which means the scope of our treatment is necessarily broad. We advocate for future studies to expand on our work.

## Methods

Our novel approach to bioacoustic source separation involves an end-to-end pipeline consisting of multiple discrete steps, including (1) synthesizing a dataset, (2) developing and training a separator network to disentangle the input mixture, and (3) constructing and training a classifier model to employ as a downstream evaluation task. This workflow requires few species-level hyperparameter modifications to account for unique vocal behavior across different biological taxa but is otherwise general and makes no assumptions about the spectrotemporal structure of the overlapping calls. We develop a complete framework for bioacoustic source separation in a permutation-invariant mode using overlapping waveforms drawn from the same class of signals. We apply BioCPPNet to macaques, dolphins, and Egyptian fruit bats, and we consider two or three concurrent “speakers”. Note that we henceforth refer to non-human animal signalers as “speakers” for consistency with the human speech separation literature^2^. We address both the closed speaker regime in which the training and evaluation data subsets contain calls produced by individuals drawn from the same distribution as well the open speaker regime in which the model is tested on calls generated by individuals not present in the training dataset.

### Bioacoustic Data

We investigate a set of species with dissimilar vocal behaviors in terms of spectral and temporal properties. We apply BioCPPNet to a macaque coo call dataset^37^ consisting of 7285 coos produced by 8 unique individuals; a bottlenose dolphin signature whistle dataset^38^ comprised of 400 signature whistles generated by 20 individuals, of which we randomly select 8 for the purposes of this study; and an Egyptian fruit bat vocalization dataset^39^ containing a heterogeneous distribution of individuals, call types, and call contexts. In the case of the bat dataset, we extract the data (31399 calls) corresponding to the 15 most heavily represented individual bats, reserving 12 individuals (27586 calls) to address the closed speaker regime and the remaining 3 individuals (3813 calls) to evaluate model performance in the open speaker scenario.

### Datasets

Motivated by WSJ0-2mix^2^, a preeminent reference dataset used for human single-channel acoustic source separation, we adopt a similar approach of constructing bioacoustic datasets by mixing multiple nonoverlapping ground truth vocalizations produced by individual animals to enable supervised training of our model. The dataset consists of an input array of the composite mixture waveforms, a target array containing the separated ground truth waveforms corresponding to the respective mixtures, and a class label array denoting the individual identities of the vocalizing animals responsible for generating the signals. For each species, the mixture dataset is generated from a corpus of bioacoustic recordings annotated according to the timestamps of the signal and the identity of the signaller. In the case of macaques, we here consider closed speaker set mixtures of two and three simultaneous speakers, but our method is functionally not limited in the number of sources (*N*) it can handle. For dolphins, we consider the closed speaker regime with two overlapping calls, and for bats, we consider both the closed and open speaker scenarios with two overlapping sources.

In general, we construct the dataset as follows. We first extract the labeled waveforms either by truncating or zero-padding the waveforms to ensure that all the samples are of fixed duration. We select the number of frames either by computing the mean plus three-sigma of the durations of the calls contained in the corpus from which we draw the signals, by selecting the maximum duration of all calls, or by choosing a fixed value. For macaques, dolphins, and bats, we use 23156 frames (0.95s), 290680 frames (3.03s), and 250000 frames (1.0s), respectively. We then randomly select vocalizations from *N* different speakers drawn from the distribution of individuals used in the study (8 macaques, 8 dolphins, 12 bats for the closed speaker regime, 3 bats for the open speaker regime) and mix them additively, ensuring to randomly shift the overlaps to simulate a more plausible scenario and to provide for asynchronicity of start times, an important acoustic cue that has been suggested as a mechanism with which the animal brain can solve the CPP^1^. Despite higher computational and memory costs, we opt to use native sampling rates, since certain animal vocalizations may reach frequencies near the native Nyquist frequency. With this in mind, however, our method does provide for resampling when the vocalizations of the particular species of interest are amenable to downsampling. Explicitly, for the three species we consider including macaques, dolphins, and bats, we use sampling rates of 24414 Hz, 96 kHz, and 250 kHz, respectively. For the closed speaker regime, the training and evaluation subsets contain calls produced by the same distribution of individuals to ensure a closed speaker set. We segment the original nonoverlapping vocalizations into 80/20 training/validation subsets. We generate the mixture training waveforms using 80% of the calls, and we construct the mixture validation subset using the remaining 20% of calls held out from the training data. In the case of overlapping bat calls (for which the corpus of bioacoustic recordings contains 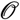(10 hours) of data as opposed to 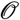(10^-1^ hours) for macaques and dolphins), we also address the open speaker source separation problem by constructing a further testing data subset of mixtures of calls of additional vocalizers not contained in the training distribution. For macaques, we construct a training data subset comprised of 12k samples and a validation subset with 3k samples, all of which contain calls drawn from 8 animals. For dolphins, we randomly select 8 individuals and construct training/validations subsets with 8k and 2k samples, respectively. For bats, we select 15 individuals, randomly reserving 12 for the closed speaker problem and the remaining 3 for the open speaker situation. We train the bat separator model on 24k mixtures. We evaluate performance in both the closed and open speaker scenarios using data subsets consisting of 6k mixtures containing unseen vocalizations produced by the appropriate distribution of individuals according to the regime under consideration. We repeat the bat training using a larger mixture dataset (denoted by +) containing 72k samples. We here report validation metrics to ensure that we are evaluating model performance on unseen mixtures of unseen calls in the closed speaker regime and on unseen mixtures of unseen calls of unseen individuals in the open speaker regime.

For the downstream classification task, we extract vocalizations annotated according to the individual identity, and we segment the calls into an 80/20 training/testing split to ensure that we are evaluating model performance on unseen calls. For both the training and evaluation data subsets, we employ an augmentation scheme in which we apply random temporal shifts to call onsets to better reflect more plausible real-world scenarios.

### Classification Models

In an effort to provide a more physically interpretable evaluation metric to supplement the commonly-implemented SI-SDR used in human speech separation studies, we develop CNN-based classifier models to label the individual identity of the separated vocalizations as a downstream task. This requires training classification networks to predict the speaker class label of the original unmixed waveforms. For each species we consider, we design and train custom simple and lightweight CNN-based architectures motivated by previous work^30^, tailored to accommodate the unique vocal behavior of the given species.

The first layer in the model is an optional high pass filter constructed using a nontrainable 1D convolution (Conv1D) layer with weights determined using a windowed sinc function to eliminate low-frequency background noise. We omit this computationally intensive layer for macaques and Egyptian fruit bats, but we implement a high pass filter for the dolphin dataset, selecting an arbitrary cutoff frequency of 4.7 kHz to remove background without impinging on the region of support for dolphin whistles. After the optional filter is an encoder layer to compute on-the-fly feature extraction. We experimented with a fully learnable free Conv1D filterbank, a spectrogram, and a log-magnitude spectrogram and observed optimal performance using a non-decibel (dB)-scaled STFT layer computed with a *nfft* window width, a *hop* window shift, and a Hann window where *nfft* and *hop* are species-dependent variables. For macaques, we select *nfft*=1024 and *hop*=64 corresponding to temporal scales on the order of 40ms and frequency resolutions on the order of 20 Hz. We choose *nfft*=1024 and *hop*=256 for dolphins and *nfft*=2048 and *hop*=512 for bats, corresponding to temporal resolutions of 10ms and 8ms and frequency resolutions of 90 Hz and 120 Hz, respectively.

Following the built-in feature engineering, the architecture includes 4 convolutional blocks, which consist of two sequential 2D convolution (Conv2D) layers with leaky ReLU activation and a max pooling layer with pool size 4. Next is a dense fully connected layer with leaky ReLU activation followed by another linear layer with log softmax activation to output the *V* log probabilities (i.e. confidences) where *V* is the number of individual vocalizers used in the study (8, 8, 12 for macaques, dolphins, and bats, respectively). We also include dropout regularization with p=0.25 for the macaque classifier and p=0.5 for the dolphin and bat classifiers to address potential overfitting. With these architectures, the macaque, dolphin, and bat classifier models have 230k, 279k, and 247k trainable parameters, respectively.

For all species, we minimize the negative log-likelihood objective loss function using the Adam optimizer^40^ with learning rate lr=3e-4. For macaques, dolphins, and bats, respectively, we train for 100, 50, and 100 epochs with batch sizes 32, 8, and 8. We save the model after each epoch and select the top-performing models. We opt not to carry out hyperparameter optimization since the classification task is of secondary importance and is used solely as a downstream task.

### Separation Models

BioCPPNet (Fig. 1) is a lightweight and modular architecture with a modifiable representation encoder, a 2D UNet core, and an inverse transform decoder, which acts directly on raw audio via on-the-fly learnable or handcrafted transforms. The structure of the network is designed to provide for extensive experimentation, optimization, and enhancement across a range of species with variable vocal behavior.

**Figure 1.**
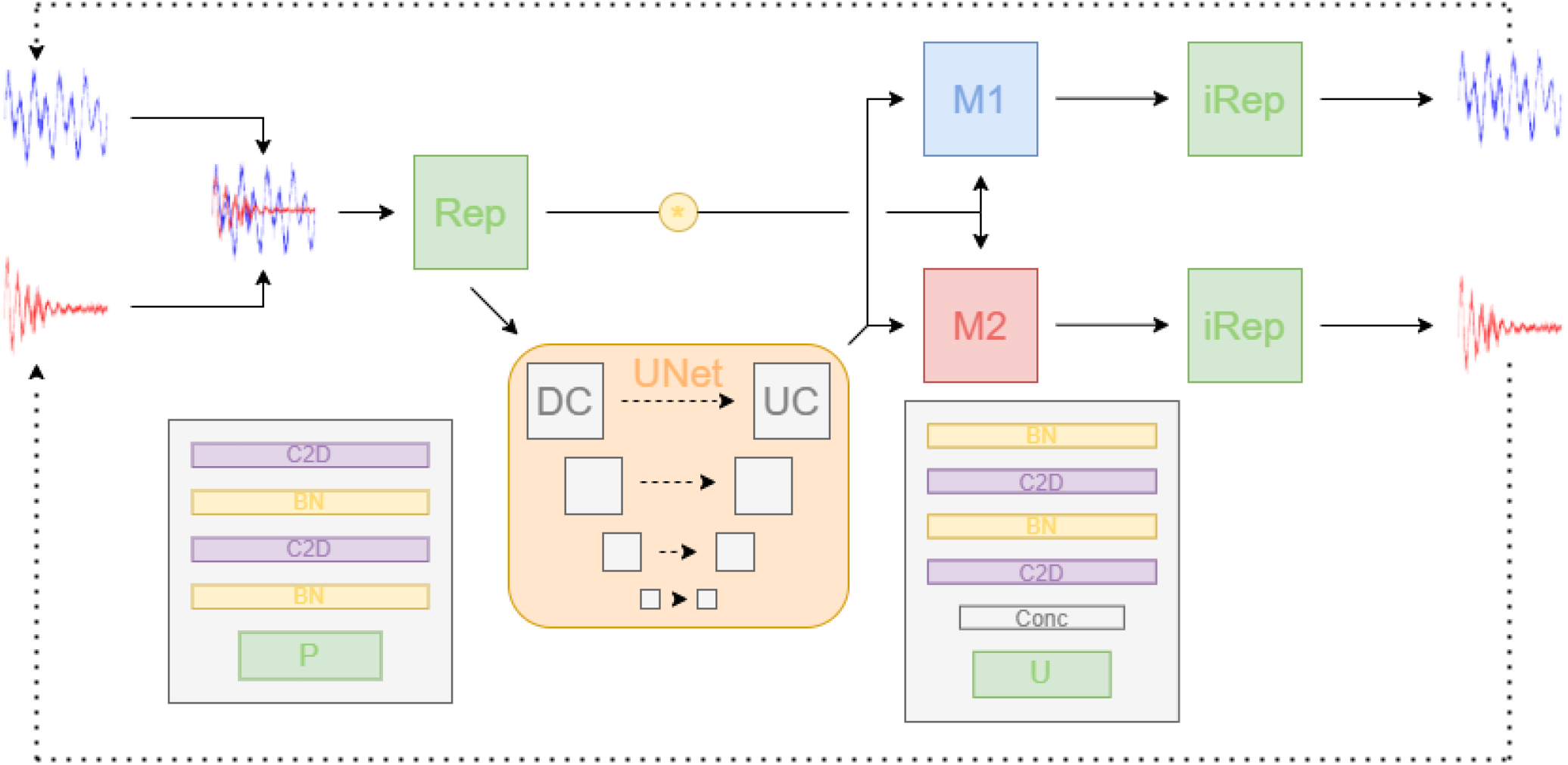
Source vocalization waveforms are mixed before being transformed to a learnable or handcrafted representation (Rep), which then passes through a 2-dimensional UNet^41^ composed of downsampling convolutional (DC) and upsampling convolutional (UC) blocks. The contracting blocks consist of two convolutional layers (C2D) with leaky ReLU activation and batch normalization (BN) followed by a max pooling (P). The expanding path consists of an upsampling (U) and skip connection concatenation (Conc) followed again by the C2D BN layers. The UNet predicts masks (M1 and M2), the number of which is determined by the number of sources (*N*), that are multiplicatively applied to the original mixture representation. The predicted representations for the separated waveforms are inverted (iRep) to output the raw waveforms, which are compared to the source ground truths, up to a permutation.

**Figure 2.**
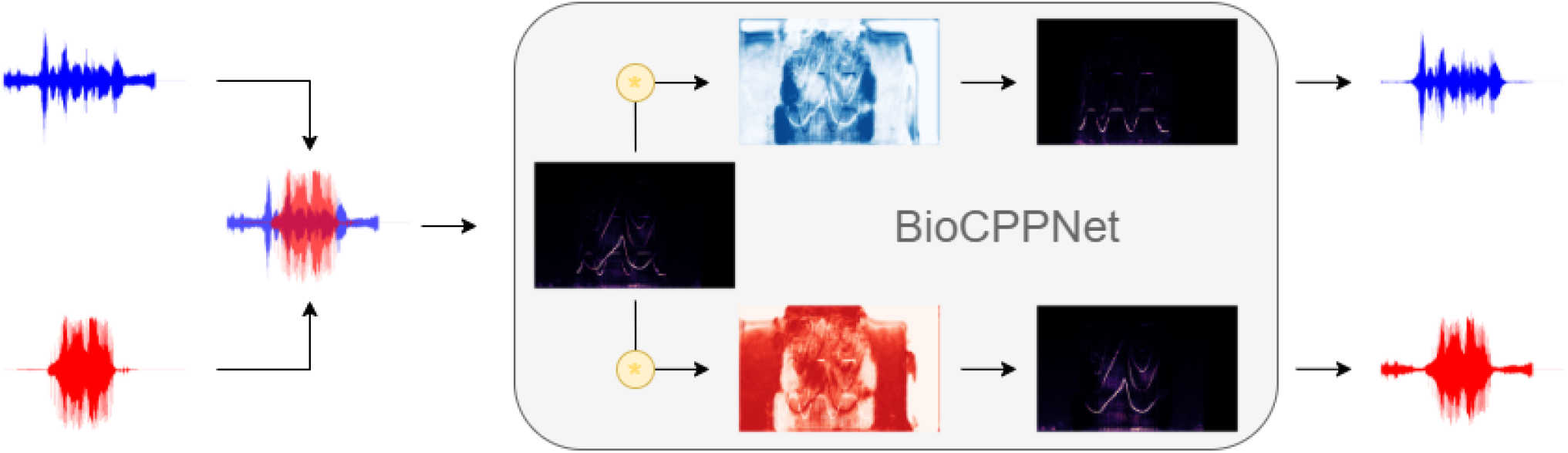
A schematic diagram demonstrating the application of BioCPPNet to dolphin signature whistles using handcrafted STFT-based encoders and decoders. The source waveforms are overlapped and transformed to time-frequency space using an STFT layer. The UNet predicts masks that are applied to the mixture representation. The estimated spectrogram separations are inverted using an iSTFT layer, and the raw output waveforms are compared with the ground truth waveforms.

#### Model Architecture

As with the classifier model, the network’s encoder consists of a feature engineering block, the initial layer of which is an optional high pass filter. This is followed by the representation transform, which includes several options including the Conv1D free encoder, the STFT filterbank, and the log-magnitude (dB) STFT filterbank. We choose the same kernel size (*nfft*) and stride (*hop*) parameters defined in the classifier model. Sequentially following the feature extraction encoder is a 2D UNet core. This architecture consists of *B* (4 for macaques, 3 for dolphins, and 4 for bats) downsampling convolutional blocks, a middle convolutional block, and *B* upsampling convolutional blocks. The downsampling blocks consist of two 2D convolutional layers with filter number that increases with model depth with leaky ReLU activation followed by a max pooling with pool size 2, 6, and 3 for macaques, dolphins, and bats. The middle block contains two 2D convolutional layers with leaky ReLU activation. The upsampling blocks include an upsampling using the bilinear algorithm and a scale factor corresponding to the pool size used during downsampling, followed by skip connections in which each corresponding level of the contracting and expanding paths are concatenated before passing through two 2D convolutional layers with leaky ReLU activation. All convolutional layers in the downsampling, middle, and upsampling blocks include batch normalization after the activation function to stabilize and expedite training and to promote regularization. Though our default implementation is phase-unaware, we also offer the option for a parallel UNet pathway working directly on phase information, which has been shown to improve performance in other applicatìons^42–44^. The final layer in the UNet core is a 2D convolutional layer with *N* channels, which are then split prior to entering the inverse transform decoder. For the inverse transform, we again provide numerous choices including a free filterbank decoder based on a 1D convolutional transpose layer, an iSTFT layer, an iSTFT layer accepting dB-scaled inputs, and a multi-head convolutional neural network (MCNN) for fast spectrogram inversion^45^. In detail, the UNet returns *N* masks that are then multiplied by the original encoded representation of the mixture waveform. The separated representations are then passed into the inverse transform layer in order to yield the raw waveforms corresponding to the model’s predictions for the separated vocalizations. We initialize all trainable weights using the Xavier uniform initialization. In the case of macaques, we experiment across all combinations of representation encoders and inverse transform decoders, and we find optimal performance using the handcrafted non-dB STFT/iSTFT layers operating in the time-frequency domain. Since the model with the fully learnable Conv1D-based encoder/decoder uniquely operates in the time domain, we report evaluation metrics for this model, as well. For dolphins and bats, we here report metrics using exclusively the non-dB STFT/iSTFT technique.

BioCPPNet (Fig. 1) is designed as a lightweight model in order to efficiently process large amounts of bioacoustic data sampled at high sampling rates while simultaneously minimizing computational costs and limitations and the likelihood of overfitting. For the macaque separators, the networks consist of 1.2M parameters (for the STFT, iSTFT combination), 2.5M parameters (for the STFT, iSTFT combination with parallel phase pathway), or 2.8M parameters (for the Conv1D free filterbanks). For the dolphin separator, the model has 304k parameters, while the bat separator model has 1.2M parameters. This is to be contrasted with the comparatively heavyweight default implementations of models commonly used in human speech separation problems, such as Conv-TasNet^3^, which has 5.1M parameters; DPTNet^4^ with 2.7M parameters; or Wavesplit^5^ with 29M parameters. Regardless of the lower complexity of BioCPPNet, the model achieves comparable performance or even outperforms reference human speech separator models while still being lightweight enough to train on a single NVIDIA P100 GPU.

#### Model Training Objective

The model training objective aims to optimize the reconstruction of separated waveforms from the aggregated composite input signal. We adopt a permutation-invariant training (PIT)^46^ scheme in which the model’s predicted outputs are compared with the ground truth sources by searching over the space of permutations of source orderings. This fundamental property of our training objective reflects that the order of estimations and their corresponding labels from a mixture waveform is not expressly germane to the task of acoustic source separation, i.e. separation is a set prediction problem independent of speaker identity ordering.

Source separation involves training a separator model *f* to reconstruct the source single-channel waveforms given a mixture 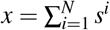 of *N* sources, where each source signal *s^i^* for *i* ∈ [1, *N*] is a real-valued continuous vector of fixed dimension *T*, i.e., *s^i^* ∈ IR^1×*T*^. The model outputs the predicted waveforms 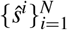 where ∀i, *ŝ^i^* = *f^i^*(*x*), and a loss function is evaluated by comparing the predictions to the ground truth sources 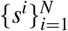 up to a permutation. Explicitly, we consider a permutation-invariant objective function^5^,

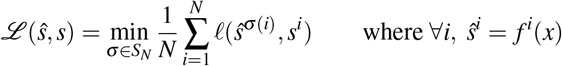

Here, *ℓ*(·, ·) represents the loss function computed on an (output, target) pair, σ indicates a permutation, and *S_N_* is the space of permutations. In certain scenarios, we include the L2 regularization term,

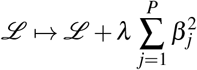

where *β_j_* represent the model parameters, *P* denotes the model complexity, and *λ* is a hyperparameter empirically selected to minimize overfitting (i.e. enhance convergence of training and evaluation losses and metrics).

We consider the reconstruction performance by computing evaluation metrics using a similar expression^5^,

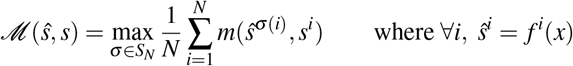

where *m*(·, ·) is the single-channel evaluation metric computed on permutations of (output, target) pairs.

For the single-channel loss function *ℓ*, we consider a linear combination of several loss terms that compute the error in estimated waveform reconstructions 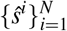 relative to the ground truth waveforms 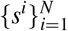.

- L1 Loss

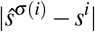 This represents the absolute error on raw time domain waveforms.
- STFT L1 Loss

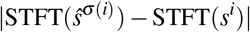 This term functions to minimize absolute error on time-frequency space representations. Empirically, the inclusion of this contribution enhances the reconstruction of signal harmonicity.
- Spectral Convergence Loss

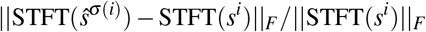

where || · ||_F_ denotes the Frobenius norm over time and frequency. This term emphasizes high-magnitude spectral components.

We also experimented with additional terms including L1 loss on log-magnitude spectrograms to address spectral valleys and negative SI-SDR (nSI-SDR), but the inclusion of these contributions did not yield empirical improvements in results.

To assess the quality of reconstruction, we compute two evaluation metrics. As is commonly implemented in the human speech separation literature, we compute the signal-to-distortion ratio (SDR)^2^, defined as the negative log squared error normalized by reference signal energy, SDR(*ŝ*, *s*) = — 10log_10_(|*ŝ* — *s*|^2^) + 10log_10_(|*s*|^2^). Further, we consider the scale-invariant SDR (SI-SDR), which disregards prediction scale and searches over gains^29^.

To provide a physically interpretable metric, we evaluate the performance of the trained classifier models in labeling separated waveforms according to the predicted identity of the vocalizer. This metric assumes that the classification accuracy on a downstream task reflects the fidelity of the estimated signal relative to the ground truth source and thus serves as a proxy for reconstruction quality.

For macaques, we modify the training algorithm according to the representation transform and inverse transform built into the model. For the model with the fully learnable Conv1D encoder and decoder, we train using the AdamW^47^ optimizer with a learning rate 3e-4 and batch size 16 for 100 epochs. In order to stabilize training and avoid local minima when using handcrafted STFT and iSTFT filterbanks, we initially begin training the models for 3 epochs with batch size 16 using stochastic gradient descent (SGD) with Nesterov momentum 0.6 and learning rate 1e-3 before switching to the AdamW optimizer until reaching 100 epochs.

For dolphins, we provide the model with the original mixture as input, but we use high pass-filtered source waveforms as the target, which means the separation model must additionally learn to denoise the input. We again initialize training with 3 epochs and batch size 8 using SGD with Nesterov momentum 0.6 and learning rate 1e-3 before switching to the AdamW optimizer with learning rate 3e-4 for the remaining 97 epochs. We use a similar training scheme for bats, initially training with SGD for 3 epochs before employing the optimizer switcher callback to switch to AdamW and to complete 100 epochs.

## Results

We here report the validation metrics for the classifier and separator models. For the closed speaker separation task, we report both the maximal SI-SDR and classification accuracies since it remains unclear which metric reflects optimal performance of the separator model. Finally, we evaluate an open speaker bat separator model using a dataset containing vocalizations produced by individuals not contained in the training distribution. The results are summarized in Table 1. Supplementary visualizations and audio samples are included in our GitHub repository https://github.com/earthspecies/cocktail-party-problem.

**Table 1.** Summary of experiments implemented in this study organized according to species, number of speakers, open/closed speaker regime, and model architecture. STFT represents the model with STFT encoder and iSTFT decoder, and Conv1D represents the fully time domain model with Conv1D encoder and ConvTranspose1D decoder. The + denotes the expanded training dataset. We report SI-SDR with (Δ) and without improvement over input SI-SDR averaged over mixtures.

| | Objective | Model | SI-SDR | $\Delta$ SI-SDR | Accuracy |
| --- | --- | --- | --- | --- | --- |
| Macaque | 2SpeakerClosed | STFT | 26.1 | 26.1 | 93.7% |
|  |  | Conv1D | 24.5 | 24.5 | 86.4% |
|  |  | Conv-TasNet | 23.8 | 23.8 | 85.1% |
|  | 3SpeakerClosed | STFT | 16.9 | 22.0 | 86.5% |
| Dolphin | 2SpeakerClosed | STFT | 15.5 | 17.3 | 99.9% |
| Bat | 2SpeakerClosed | STFT | 9.7 | 9.7 | 57.5% |
|  | 2SpeakerOpen |  | 10.0 | 10.0 | — |
|  | 2SpeakerClosed+ |  | 10.3 | 10.3 | 58.6% |
|  | 2SpeakerOpen+ |  | 10.4 | 10.4 | — |

### Classification Models

The macaque classifier model attains an accuracy of 99.3% (where chance level was 12.5%, i.e. 1 out of 8) for the 8 vocalizing animals used in this study, which represents a state-of-the-art improvement over the existing literature^16^. The dolphin classifier model achieves 99.4% accuracy (where chance level was 12.5%, i.e. 1 out of 8) in classifying signature whistles produced by 8 individuals, again reinforcing the individually distinctive characteristics of dolphin signature whistles^38^. The Egyptian fruit bat classifier network yields 79.7% accuracy (where chance level was 8.3%, i.e. 1 out of 12) in labeling the identity of 12 bats, supporting previous studies demonstrating individual-level specificity of everyday bat vocalizations^48^.

### Separation Models

For the macaque separation task, we here detail the results for the fully learnable Conv1D-based model as well as the top-performing non-dB STFT-based model. We also consider mixtures of 2 or 3 overlapping speakers. For the baseline model using the fully learnable Conv1D encoder/decoder, the model yields an SI-SDR of 24.5 and a downstream classification accuracy of 86.4%. We experiment across combinations of (Conv1D, STFT, dB-scaled STFT) encoders and (Conv1D, iSTFT, dB-scaled iSTFT, MCNN) decoders. We observe that the handcrafted (STFT, iSTFT) encoder/decoder combination results in the highest fidelity reconstructions, as assessed using SI-SDR and downstream classification accuracy, with respective (SI-SDR, accuracy) metrics of (26.1, 93.7%). Including the parallel phase pathway does not yield robust performance benefits, in contrast with results from other experimental studies^42^. We note that, even though our model is significantly less complex and computationally expensive, it outperforms Conv-TasNet, a pioneering human speech separation model, which attains a maximal evaluation metric pair of (23.8, 85.1%). Finally, training an STFT/iSTFT model with *N*=3 yields evaluation metrics of (16.9, 86.5%). For both the dolphin separation and bat separation tasks, we report the results using exclusively the fixed STFT encoder and iSTFT decoder rather than carrying out an ablation study across various TFRs and inverses. The dolphin separator yields metrics of (15.5, 99.9%) and the bat model achieves (9.7, 57.5%). Finally, to address the open speaker bat source separation problem, we evaluate the bat model on (1) the training subset, (2) the closed speaker testing subset, and (3) the open speaker testing subset, yielding respective SI-SDR metrics of 10.5, 9.7, 10.0. This finding demonstrates that even in the absence of individually recognizable acoustic cues, BioCPPNet generalizes to overlapping waveforms produced by unseen vocalizers not contained in the training subset. We repeat the bat source separation problem using the larger 72k-sample training dataset (+) and observe maximal metrics of (10.3, 58.6%) in the closed speaker regime. For the open speaker regime, the respective training, closed speaker testing, and open speaker testing SI-SDRs are 10.2, 10.3, 10.4, indicating that training on more samples improves bat separator model generalizability. In Table 1, we include the SI-SDR with improvement (Δ SI-SDR), i.e. the metric obtained using the model output minus the metric averaged over input mixtures.

## Discussion

This work proposes BioCPPNet, a complete supervised end-to-end bioacoustic source separation framework designed to accommodate multiple species with diverse vocal behavior. We apply our model to a heterogeneous set of non-human organisms including macaques, bottlenose dolphins, and Egyptian fruit bats, and we demonstrate that our models yield high-quality reconstructions of sources given mixture inputs, as assessed objectively using the SI-SDR metric in combination with downstream classification accuracy and qualitatively by inspecting visual representations of the output audio. Though BioCPPNet performs best in the closed speaker regime in which testing subsets are drawn from the same distribution of individuals as the training subset, our results suggest that BioCPPNet can generalize to the open speaker problem given sufficient quantities of data; for instance, we observe that BioCPPNet and the Conv-TasNet reference model struggle in the open speaker regime when tested on macaques (for which we possess fewer than 45 minutes of highly stereotyped calls) and bottlenose dolphins (for which the dataset is comprised of fewer than 15 minutes of signature whistles). However, when evaluated on bat data containing several hours of varied bat vocalizations, our model yields comparable results in both the open and closed speaker cases. This finding highlights the need for larger datasets to facilitate the development of novel ML-based technologies for bioacoustic source separation applications.

Given the flexibility, versatility, and modularity of BioCPPNet, we suggest further experimental setups to expand on our results. While we implement our methods using both fully learnable and STFT-based handcrafted encoder and decoder filterbanks, future directions can include modifications to the TFR encoder and the inverse transform decoder to assess the performance of other learnable, handcrafted, and/or parameterized features. To our knowledge, there exists no landscape analysis of TFRs for bioacoustic data, so additional studies aiming to enhance separation performance by varying the representation encoder and the inverse transform decoder could provide important insight into optimizing the representation of bioacoustic signals. Additionally, though we implement a learnable signal filtering mechanism in the case of bottlenose dolphin whistles, our results could benefit from additional denoising or enhancement algorithms^49–52^; as in Fig. 3 (a) and (b), noisy artifact is not always attributed to the original source, which means that targeting and removing noise in the spectral zone of support for a given set of vocalizations could improve the separation performance of BioCPPNet. We also suggest implementations of BioCPPNet across other non-human animals in an effort to further examine the generalizability of our approach.

**Figure 3.**
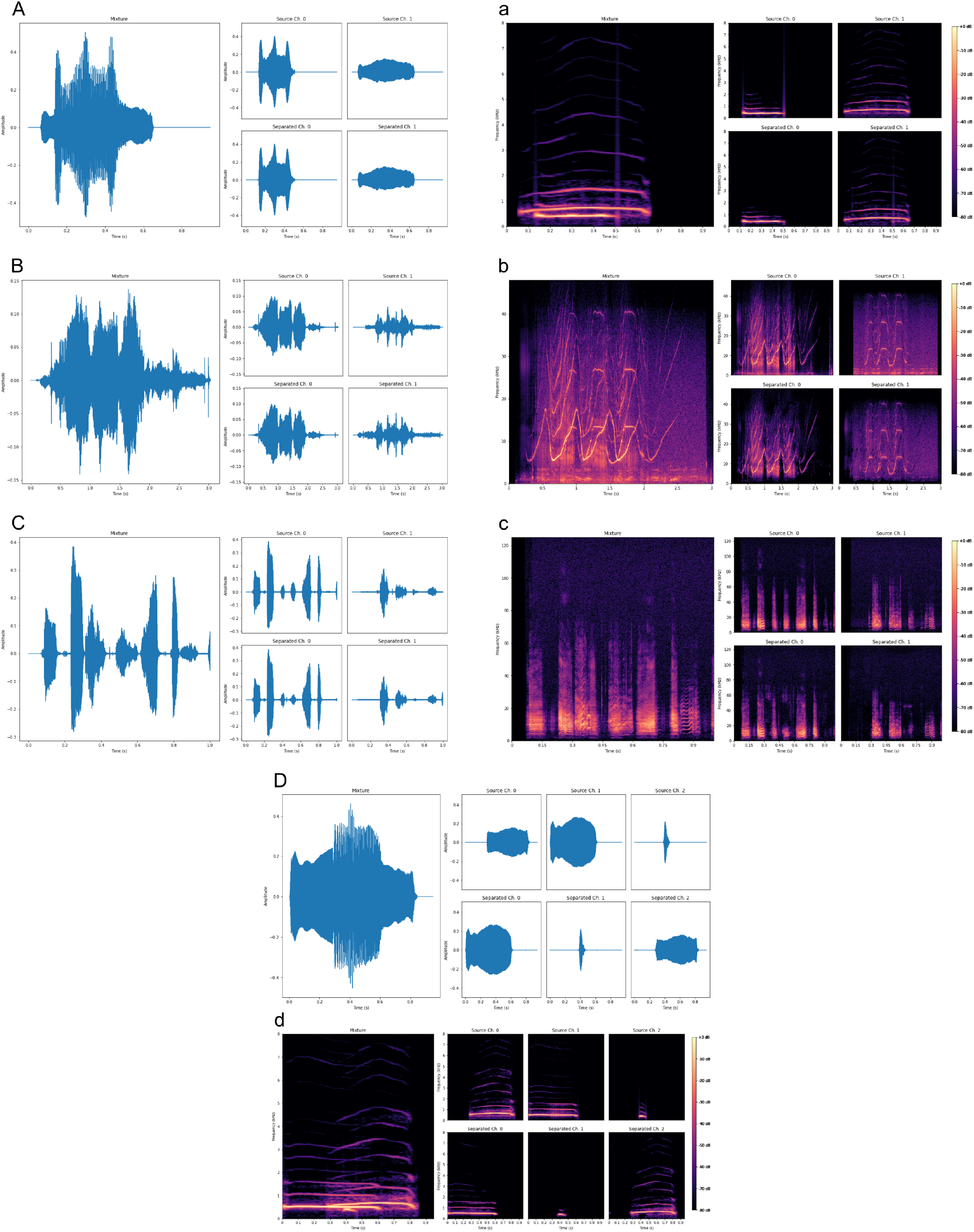
Visualizations of the mixture separations for (A,a) macaques, (B,b) bottlenose dolphins, and (C, c) Egyptian fruit bats. (D,d) demonstrates the separation of three overlapping macaque vocalizations.

Finally, while our implementation uses a supervised training scheme relying on synthesized mixtures and access to ground truth sources, the practical application of our methods could benefit from weakening the degree of supervision. Mixture invariant training (MixIT)^53^ represents an entirely unsupervised approach and requires only single-channel acoustic mixtures, though this technique may be limited in the bioacoustic domain as it necessitates large quantities of well-defined mixtures. Another unsupervised technique includes a Bayesian approach employing deep generative priors^54–56^, but this method may be limited to data within close bounds of the training sets since the distributions learned by acoustic deep generative models may not exhibit the right properties for probabilistic source separation^57^. We also suggest self-supervised pre-training^30,58,59^ on relevant proxy tasks to enhance performance, especially in the low-data bioacoustic domain. Future studies should address unsupervised training criteria in an effort to apply deep ML-based models such as BioCPPNet to real-world bioacoustic mixtures.

The novel ML-based methods employed in this study provide an effective means for addressing the bioacoustic CPP, which can help to maximize the information extracted from bioacoustic recordings. Our framework represents an important step toward the realization of bioacoustic processing technologies capable of recovering large quantities of previously unusable data containing overlapping signals. Greater access to larger bioacoustic datasets can empower the scientific community to explore more areas of research, to confront increasingly complex research questions using deep ML approaches, and ultimately to design better-informed conservation and management strategies to protect the earth’s non-human animal species.

## Data Availability

The macaque coo dataset^37^ that supports the findings of this study is publicly available in *Dryad* with the identifier https://doi.org/10.5061/dryad.7f4p9, and the Egyptian fruit bat dataset^39^ used in this study is publicly available in *Scientific Data* with the identifier https://doi.org/10.1038/sdata.2017.143. The bottlenose dolphin whistle data that support the findings of this study are available from Laela Sayigh but restrictions apply to the availability of these data, which were used under license for the current study, and so are not publicly available. Data are however available from the author upon reasonable request and with permission of Laela Sayigh.

## Code Availability

The custom-written code for generating mixture datasets and constructing and training ML models is publicly available at our GitHub repository https://github.com/earthspecies/cocktail-party-problem.

## Acknowledgements

The author thanks Aza Raskin, Britt Selvitelle, and Katherine Zacarian for useful discussion and Laela Sayigh, Frants Jensen, Michelle Fournet, Andrés Babino, and James Crutchfield for reviewing the manuscript.

## Author Information

### Contributions

All aspects of the work (task setting, data processing, machine learning, article writing, etc.) are performed by the author.

